# Dihydropyrimidinase-like 2 (DPYSL2) regulates breast cancer migration via a JAK/STAT3/vimentin axis

**DOI:** 10.1101/2021.03.21.436294

**Authors:** Areej Abu Rmaileh, Balakrishnan Solaimuthu, Michal Ben Yosef, Anees Khatib, Michal Lichtenstein, Mayur Tanna, Arata Hayashi, Nir Pillar, Yoav D. Shaul

## Abstract

The intricate neuronal wiring during development requires cytoskeletal reorganization orchestrated by signaling cues. Considering that cytoskeletal remodeling is a hallmark of cell migration, we inquired whether metastatic cancer cells exploit the axon guidance proteins to migrate. Indeed, in breast cancer patients, we found a significant correlation between the mesenchymal markers and the expression of dihydropyrimidinase-like 2 (DPYSL2), a regulator of cytoskeletal dynamics in growing axons. Strikingly, DPYSL2 knockout in mesenchymal-like cells profoundly inhibited cell migration, invasion, stemness features, tumor growth rate, and metastasis. Next, we aimed to decode the molecular mechanism underlying this phenomenon and revealed an interaction between DPYSL2 and Janus kinase 1 (JAK1). This binding is crucial for triggering signal transducer and activator of transcription 3 (STAT3) and subsequently expressing vimentin, the pro-migratory intermediate filament. Collectively, we identified DPYSL2 as a molecular link between oncogenic signaling pathways and cytoskeletal reorganization in migrating breast cancer cells.

**Statement of significance:** This study shows that the axon guidance adaptor protein DPYSL2 is essential for promoting breast cancer migration. Specifically, this protein interacts with JAK1 to govern STAT3 signaling and subsequently vimentin expression.

## Introduction

The guidance of developing neurons to their specific target region is regulated by a combination of signaling cues composed of ligand/receptor interactions (1). The axon guidance machinery includes distinct families of canonical guidance proteins, such as slit guidance ligand (SLIT) (2). This family of secreted factors induces changes in the newly developing axon through interaction with members of the roundabout guidance receptor (ROBO) family (3). Other axon guidance families include the semaphorins and their receptors, neuropilins (NRP) (4). The outcome of semaphorins/NRP and ROBO/SLIT activation is structural reorganization of the cell (5) mediated by cytoskeleton binding proteins (6), such as the collapsin response mediator protein (CRMP) family (7). Thus, in development cytoskeletal rearrangement is a central process, which is necessary for the axons to navigate to their defined targets.

Epithelial-mesenchymal transition (EMT) was initially described as an early developmental program used by epithelial cells to trans-differentiate and gain mesenchymal-like properties (8). In tumors, this program is assumed to be the mechanism by which carcinomas gain aggressive features such as migratory capabilities, chemoresistance, and stemness (9-11). The activation of the EMT program by extracellular cues (12), such as the transforming growth factor-β (TGFβ) (13,14) and interleukin 6 (IL-6) (9), results in significant transcriptome changes. Among the differentially expressed genes are cytoskeletal elements that drive the carcinoma cells to gain mesenchymal-like morphology and migratory capacity (8). Mesenchymal cell migration involves several steps that are governed by the three cytoskeletal networks, actin filaments, microtubule, and intermediate filaments (IF) (15). One of the main IFs is vimentin, an established EMT marker (16), which serves as a key regulator of cancer cell migration (17) and aggressiveness (18). Vimentin expression is induced by signaling cascades such as signal transducer and activator of transcription 3 (STAT3) (19) while its activity is regulated by posttranslational modifications including phosphorylation (20), glycosylation (21), and ubiquitination (22).

During development, the growing axon demonstrates cellular properties shared with those associated with cancer cell migration (23,24), including extracellular matrix (ECM) remodeling using specific matrix metalloproteins (MMP) (25). These MMPs include the gelatinases, MMP2 and MMP9 (26), which are established targets of the EMT program (27). The similarity between neuronal cell development and cancer cell migration prompted us to systematically characterize the expression of axon guidance genes in tumors. We identified dihydropyrimidinase-like 2 (DPYSL2, also known as CRMP2) to be highly expressed in mesenchymal-like breast cancer cells and plays an essential role in their migration ability. In these cells, DPYSL2 interacts with the Janus kinase 1 (JAK1) and mediates the JAK/STAT3 signaling, subsequently regulating the expression of the downstream target, vimentin.

## Materials and methods

### Animal Studies

MDA-MB-231 WT and DPYSL2-KO cells were injected into the mammary fat pad of female NOD-SCID mice (1 × 10^6^ cells per mouse). The tumors were monitored and measured weekly. After 6 weeks, the tumors were harvested and weighed. The obtained lungs were observed under SMZ18 Nikon Stereo microscope. The pictures were slightly edited (brightness) with Adobe Photoshop. All mouse experiments were carried out under The Hebrew University Institutional Animal Care and Use Committee-approved (IACUC) protocol MD-16-14939-5. The Hebrew University is certified by the Association for Assessment and Accreditation of Laboratory Animal Care (AAALAC)

### Immunoprecipitation

Cells were rinsed twice with ice-cold PBS and lysed in ice-cold lysis buffer (50 mM HEPES-KOH pH 7.4, 2 mM EDTA, 10 mM pyrophosphate, and 1% NP40 alternative, in addition to 0.5 mM sodium orthovanadate, 16mM sodium fluoride and one tablet of EDTA-free protease inhibitors (Roche) per 50 ml). The soluble fractions of cell lysates were isolated by centrifugation (13,000 rpm for 10 min) in a microfuge. Flag M2 affinity resins (Sigma-Aldrich) were washed with lysis buffer three times. 30 µl of a 50% slurry of the resins was then added to the cleared cell lysates and incubated with rotation for 2 h at 4°C. Finally, the beads were washed three times with lysis buffer containing 150 mM NaCl. For elution of FLAG-tagged proteins, the beads were incubated in elution buffer (50 mM HEPES-KOH pH7.4, 2 mM EDTA, 10 mM pyrophosphate, 150 mM NaCl, 1% NP40 alternative and 100 µg/ml FLAG peptide) for 1 h at 30°C with shaking. Immunoprecipitated proteins were denatured by the addition of 150 µl of SDS sample buffer (5X) and boiling for 5 min, resolved by 10% SDS-PAGE, and analyzed by immunoblotting as described.

### CRISPR/Cas9-Mediated Knockout Cell lines

We used CRISPR-Cas9-mediated genome editing to achieve gene knockout, using pLentiCRISPR v1 (Addgene Plasmid #70662) in which the sgRNA and Cas9 are delivered on a single plasmid. We also generated knockout cells using pSpCas9(BB)-2A-Puro (PX459) V2.0 (Addgene Plasmid #62988). Editing of the DPYSL2 locus in MDA-MB-231 cells was accomplished by either infecting cells with the “pLentiCRISPR” plasmid, or by transfection with the PX459 plasmid into which a sgRNA targeting the DPYSL2 locus had been cloned. Cells were then subjected to single-cell cloning by limiting dilution in 96-well plates. Editing of the DPYSL2 locus was confirmed by assessing protein level by Western Blot.

Guide RNA sequences

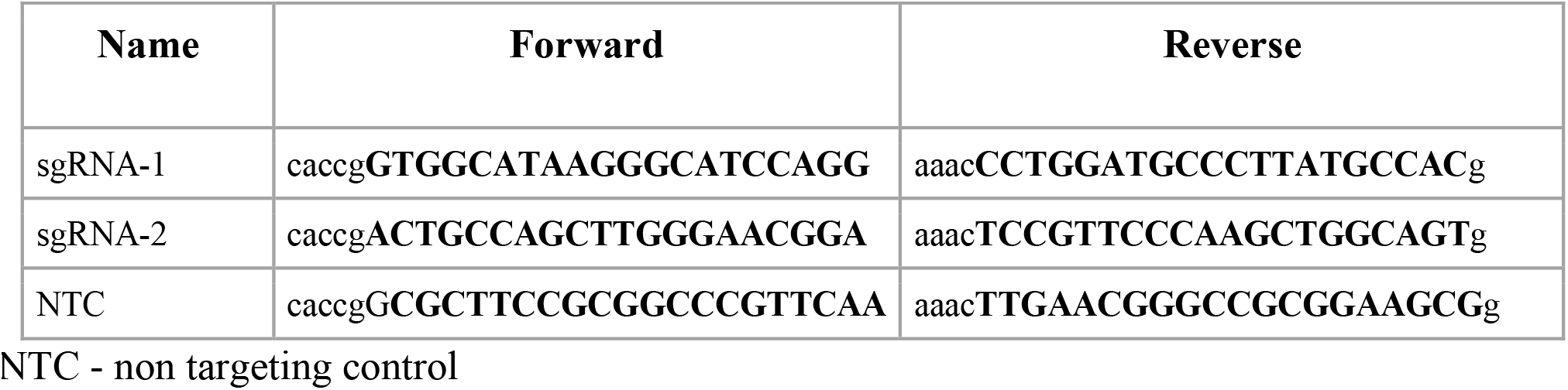

### Statistical Analysis

All statistical analyses were performed using the R (version 4.0) or GraphPad Prism (version 8.0) statistical analysis programs. If not indicated otherwise, all the *p-*values in the figures measured between the indicated samples were quantified using Student’s *t*-test.

### Analysis of breast cancer data in cBioportal

The cBioPortal for Cancer Genomics is an open-access database, providing visualization and analysis tools for large-scale cancer genomics data sets (http://cbioportal.org). For gene correlation analysis we queried DPYSL2 or JAK1 in breast invasive carcinoma (TCGA, PanCancer Atlas project) (34), or the METABRIC (39) (containing 1,084 or 2,509 samples respectively). Then, we subjected the genes to co-expression analysis and downloaded the correlation plots. For gene set enrichment analysis (GSEA), Spearman’s correlation coefficient between the gene of interest and the whole genome were computed, downloaded, and subjected to GSEA using the GSEA 4.1.0 version software (37). For the different analysis we selected the h.all.v7.2.symbole.gmt (Hallmarks) or C2.cp.kegg.v7.2.symbols.gmt (Curated) gene set databases.

## Results

### DPYSL2 expression correlates with mesenchymal markers

We sought to identify the axon guidance molecules that are differentially expressed in mesenchymal and epithelial samples. Previously, we segregated the cancer cell lines of the MERAV database into epithelial and mesenchymal groups (28) according to their transcriptome (http://merav.wi.mit.edu/) (29). We then analyzed these two groups for their expression of 126 axon guidance genes, as defined by the Kyoto encyclopedia of genes and genomes (KEGG) database (https://www.genome.jp/dbget-bin/www_bget?pathway:hsa04360). We found ten axon guidance genes that were significantly upregulated (cut off ratio of Log_2_=1, Mesenchymal-UP) and five downregulated (cut off ratio of Log_2_=-1, Mesenchymal-Down) in mesenchymal cell lines (Figure S1A). The upregulated set included genes known to participate in cancer cell aggressiveness, including *SLIT2, ROBO1* (30), *NRP1* (31), FYN proto-oncogene (*FYN*) (32), and cofilin 2 (*CFL2*) (33) (Figure 1A and B), supporting our bioinformatics analysis. Since *DPYSL2* exhibited the highest expression level in mesenchymal cells, and its function in cancer is not yet established, we decided to characterize its role in cancer cell aggressiveness.

**Figure 1:**
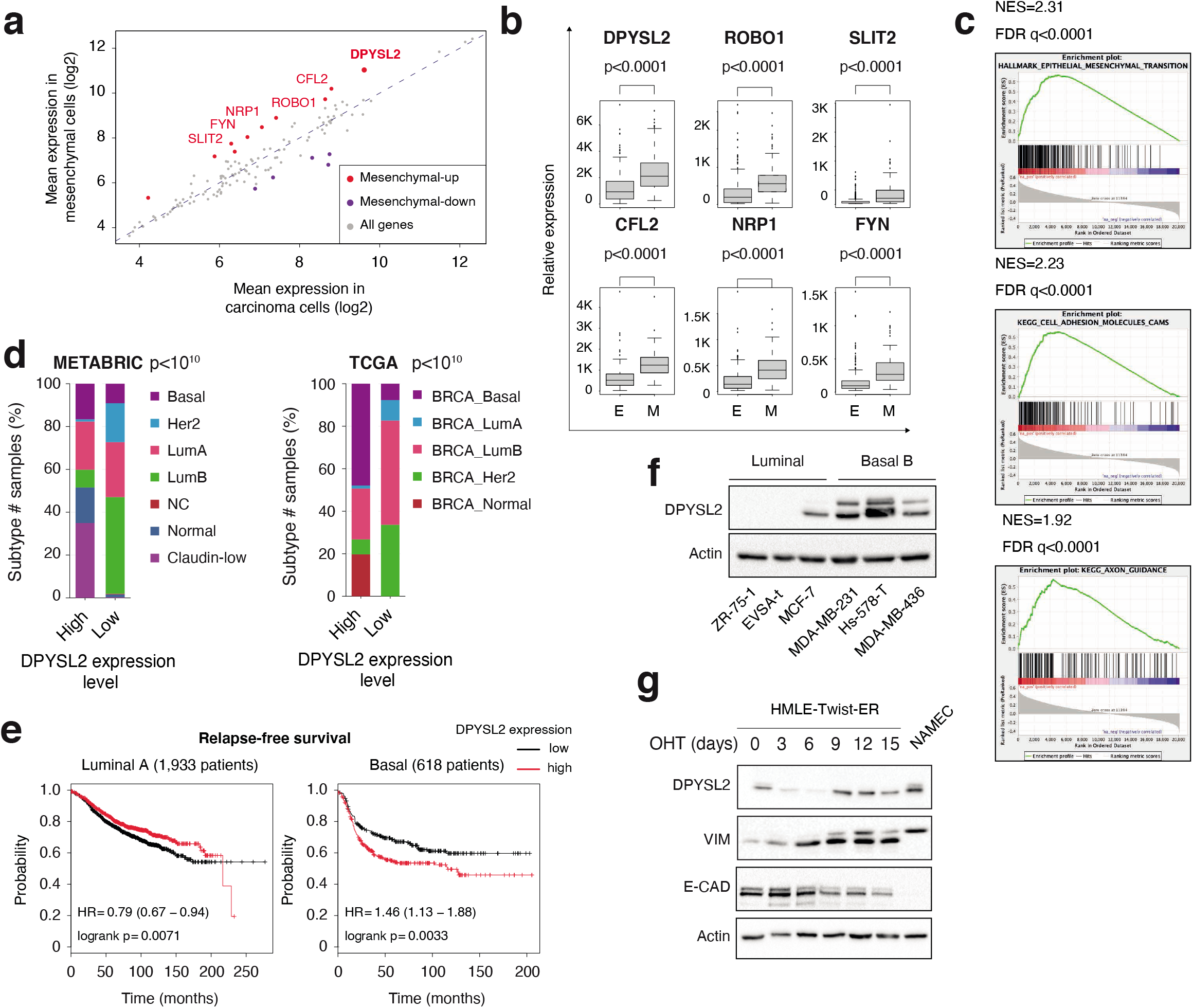
DPYSL2 expression is elevated in mesenchymal-like cells. (**a**) Several of the axon guidance genes demonstrate increased expression in mesenchymal cells. Cancer cell lines were divided into epithelial (n=378 cell lines) and mesenchymal (n=150 cell lines) groups based on the expression of known mesenchymal markers. The expression of the 126 axon guidance genes was compared to the mean expression in each group. Gray -axon guidance genes (all genes); red -genes up regulated in mesenchymal cells; purple – genes up regulated in epithelial cells. (**b**) The expression levels of selected mesenchymal-UP axon guidance genes in mesenchymal cell lines. Box plots represent the expression levels of the indicated genes in each group (as in A). The p-value was determined by Student’s t-test. (**c**) DPYSL2 expression in breast cancer patients correlates with the hallmark of EMT. Breast cancer patients’ gene expression data were generated by the TCGA (PanCancer Atlas project) and analyzed using the cBioportal webtool (https://www.cbioportal.org). In these samples, the expression of DPYSL2 was compared to the whole transcriptome (∼20,000 genes). The genes were ranked based on the obtained Spearman correlation coeffcient following by gene set enrichment analysis (GSEA). GSEA computed the normalized enrichment score (NES) and FDR values. (**d**) DPYSL2 expression is higher in aggressive breast cancer subtypes. Breast cancer samples were divided into two groups based on high and low DPYSL2 expression (one standard deviation above or below the mean). For each group, the percentage of breast cancer subtypes is color-coded. The breast cancer data was obtained from the METABRIC (left) or the TCGA (PanCancer Atlas project; right) databases, and the p-value was calculated by the cbioportal. Lum=luminal. (**e**) DPYSL2 expression is associated with poor relapse-free survival in basal cancers. Kaplan-Meier survival plots for patients with breast cancer were divided into high DPYSL2 expression (“high” (red)) and low (“low” (black)). The numbers in parentheses indicate the total number of patients. These plots were generated in the Kaplan-Meier plotter website. The DPYSL2 (200762_at Affymetrix ID symbol) was used for all the analyses. The p-value (P), and the hazard ratio (HR) were determined by the analysis tool. (**f**) The DPYSL2 protein level is upregulated in high-grade breast cancer cell lines. Cells were lysed and subjected to immunoblotting using the indicated antibodies. (**g**) DPYSL2 expression is upregulated during the EMT program. HMLE-Twist-ER cells were treated with hydroxytamoxifen (OHT) to induce EMT for a total of 15 days. Every three days, cells were collected, lysed, and subjected to immunoblotting using the indicated antibodies. NAMEC -Naturally Arising MEsenchymal Cells, an HMLE-derived cell line that spontaneously acquired the mesenchymal state.

We then assessed the DPYSL2 expression profile in patient-derived breast cancer samples. We analyzed the cancer genome atlas (TCGA, PanCancer Atlas project) data (34) available on cBioportal (https://www.cbioportal.org)(35,36), and found by gene set enrich analysis (GSEA) (37) that *DPYSL2* expression significantly correlated with EMT, cell adhesion, and axon guidance molecules (Figure 1C). We found that in breast cancer, DPYSL2 significantly correlates with the axon guidance molecules *ROBO, SLIT2, FYN*, and *NRP1* (Figure S1B). Moreover, it also significantly correlated with the EMT markers Snail Family Transcriptional Repressor 2 (*SNAI2*), zinc finger E-box binding homeobox 1, 2 (*ZEB1, ZEB2*), and vimentin (*VIM*) that demonstrated the highest Spearman’s correlation coefficient (0.61) (Figure S1C). Whereas DPYSL2 expression anticorrelated with the epithelial markers keratin 18 (*KRT18*) and keratin 19 (*KRT19*) (38) (Figure S1D). Furthermore, by analyzing both the METABRIC (39) and TCGA databases (40), we found that samples highly expressing DPYSL2 (DPYSL2-high) are significantly enriched with high-grade breast cancer subtypes (basal and claudin-low), in comparison to DPYSL2-low samples (Figure 1D). Additionally, utilizing the Kaplan-Meier Plotter tool (http://kmplot.com/analysis/) (41), we identified a significant association between high DPYSL2 expression levels and poor relapse-free survival (RFS) rate (42) (Figure 1E).

Next, we determined that DPYSL2 is upregulated in breast cancer-derived mesenchymal-like (basal B) cell lines relative to epithelial (luminal) cell lines (Figure 1F). Likewise, EMT induction in engineered human mammary epithelial (HMLE-Twist-ER) cells induced DPYSL2 expression. In this system, the EMT program is induced by 4-hydroxytamoxifen (OHT), which in turn, activates an ectopically expressed twist family BHLH transcription factor 1 (Twist1) conjugated to the estrogen receptor (28,43) (Figure 1G). This treatment also resulted in upregulation of vimentin (VIM) and downregulation of the epithelial marker, E-cadherin (E-CAD). Thus, we revealed that the EMT program regulates DPYSL2 expression, indicating its role in cancer cell aggressiveness.

### DPYSL2 loss inhibits cell aggressiveness in breast cancer cell lines

To further assess DPYSL2’s contribution to cancer aggressiveness, we knocked out DPYSL2 from the basal B breast cancer cell line, MDA-MB-231 (DPYSL2-KO), using the Cas9-CRISPR system (Figure 2A). These cells were then subjected to comparative CEL-Seq analysis, which generated a list of significantly downregulated genes in the knockout relative to WT cells (Table S1). Based on Metascape analysis (https://metascape.org/) (44), we systematically categorized this gene list according to biological functions related to migration and cellular signaling (Figure 2B).

**Figure 2:**
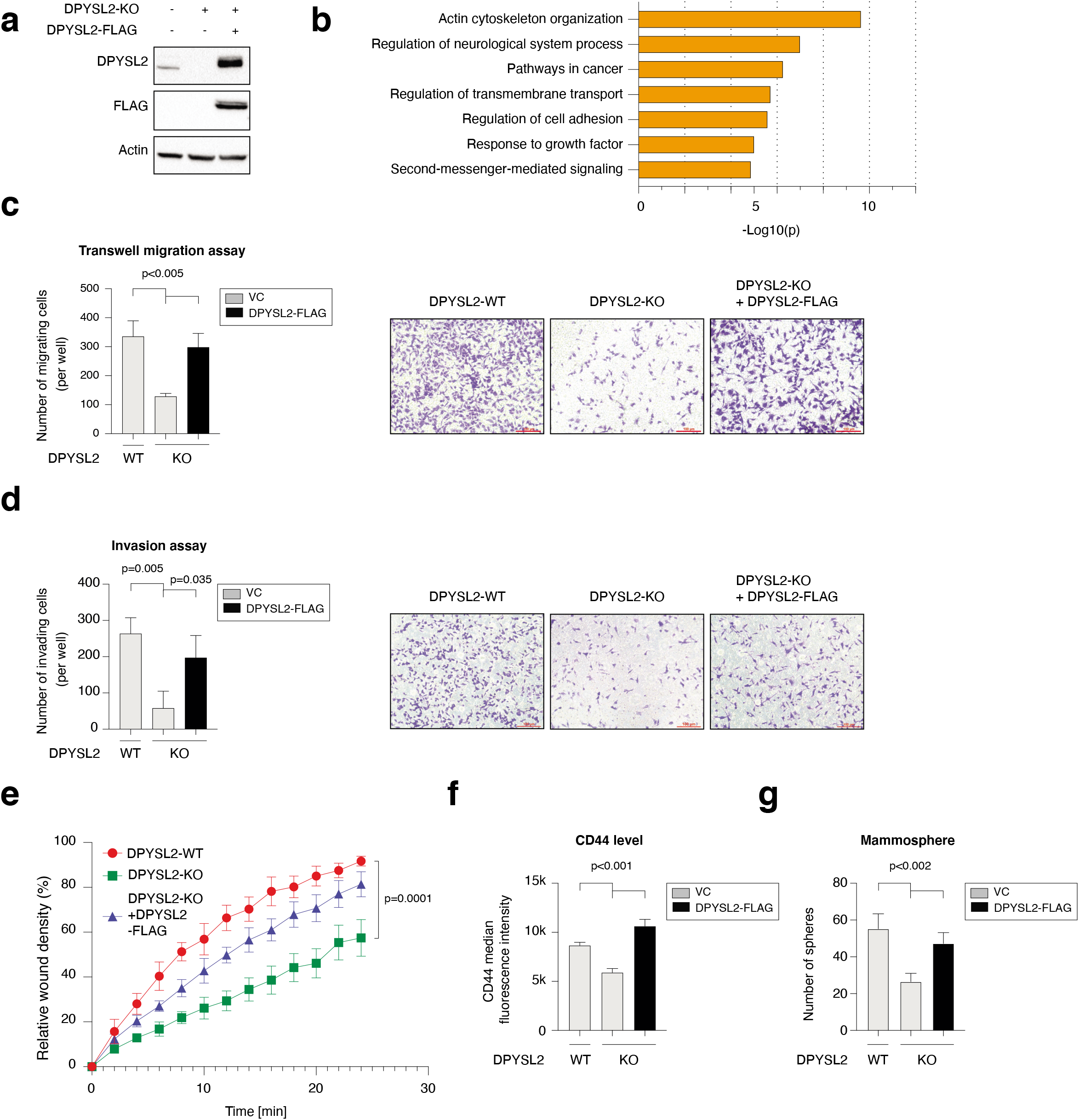
DPYSL2 loss inhibits cell aggressiveness in the breast cancer cell line MDA-MB-231. (**a**) Immunoblot representing DPYSL2 knockout (DPYSL2-KO) in MDA-MB-231 cells. FLAG-tagged DPYSL2 (DPYSL2-FLAG) was introduced into DPYSL2-KO cells. Cells were lysed and subjected to immunoblotting using the indicated antibodies. (**b**) DPYSL2 loss affects actin filaments’ remodeling and cellular signaling. Both WT and DPYSL2-KO cells were subjected to comparative CEL-Seq analysis and the expression level of each gene was analyzed in the two samples. The set of genes that demonstrated a significant reduction upon DPYSL2 loss were further subjected to Metascape analysis (https://metascape.org/). (**c**) DPYSL2 loss inhibits cell migration in breast cancer cell line. The migratory capability of the different samples was determined in a transwell assay. Quantification of data is reported as the number of migrated cells per 20,000 seeded cells (left); each value represents the mean ± SD for n=3. The p-value was determined by Student’s t-test. Representative images of each sample (right). VC -vector control. (**d**) DPYSL2 overexpression rescues the DPYSL2-KO effect on cell invasiveness. MDA-MB-231 cells were infected and treated as in (c), and the number of the matrigel-invading cells was measured. Quantification of data is reported as the number of migrated cells per 20,000 seeded cells (left); each value represents the mean ± SD for n=3. The p-value was determined by Student’s t-test. Representative images of each sample (right). VC -vector control. (**e**) Quantification of relative wound density over 24 h for the indicated WT, DPYSL2-KO, and DPYSL2-KO+ DPYSL2-FLAG cells. Each value represents the mean ± SD for n=8. (**f**) Loss of DPYSL2 results in CD44 expression reduction. The different indicated samples were subjected to FAGS analysis of the cell-surface marker GD44. The plot represents GD44 fuorescence intensity values. n=3. VC -vector control. (**g**) DPYSL2 overexpression rescues mammosphere-forming ability in DPYSL2-KO cells. A quantification of in vitro mammosphere formation by cells from the different samples was performed. The data is reported as the number of mammospheres formed per 800 seeded cells; each value represents the mean ± SD for n = 4. The p-value was determined by Student’s t-test. VG -vector control.

Next, we applied *in vitro* functional readouts to assess the involvement of DPYSL2 in cancer cell aggressiveness. Using the Boyden chamber-based transwell migration assay (Figure 2C), invasion assay (Figure 2D), and the Incucyte Live-Cell analysis system (Figure 2E and S2A), we found that DPYSL2 loss significantly inhibited the migratory and invasive abilities of these cells without affecting the proliferation rate (Figure S2B). Additionally, we found that DPYSL2-KO cells, relative to WT, showed significant reductions in the extracellular CD44 levels (Figure 2F and S2C) and mammosphere formation ability (Figure 2G), which are well-established stemness indicators (45). Furthermore, restoring DPYSL2 expression in these KO cells (DPYSL2-KO+DPYSL2-FLAG) (Figure 2A) improved their migration, invasion, and stemness to levels comparable to WT (DPYSL2-WT) (Figure 2C, D, E, F, and G). Together these *in vitro* assays indicated the crucial role of DPYSL2 in maintaining the aggressive state of cancer cells.

### DPYSL2 expression promotes tumor formation and metastasis

We aimed to determine the role of DPYSL2 in tumor formation and metastasis *in vivo*. Accordingly, we injected GFP-labeled MDA-MB-231 cells (WT or DPYSL2-KO) into the mammary fat pads of female NOD-SCID mice and monitored the tumor growth for six weeks. We found that tumor growth rate in mice injected with the DPYSL2-KO cells was significantly lower than that in mice injected with WT cells (Figure 3A). Correspondingly, the average weight of tumors formed in the DPYSL2-KO-injected mice was significantly lower than those from WT-injected mice (Figure 3B and S3A). We also harvested the lungs of the injected mice, examined them for the presence of metastases, and measured the number of GFP-positive colonies. In comparison to WT, DPYSL2-KO-injeced mice had a significant reduction in the number of lung metastases (Figure 3C, 3D, and S3B), which was further validated by hematoxylin and eosin (H&E) staining (Figure 3E and S3C). Altogether, we found that DPYSL2 is a central player in tumor growth and metastasis.

**Figure 3:**
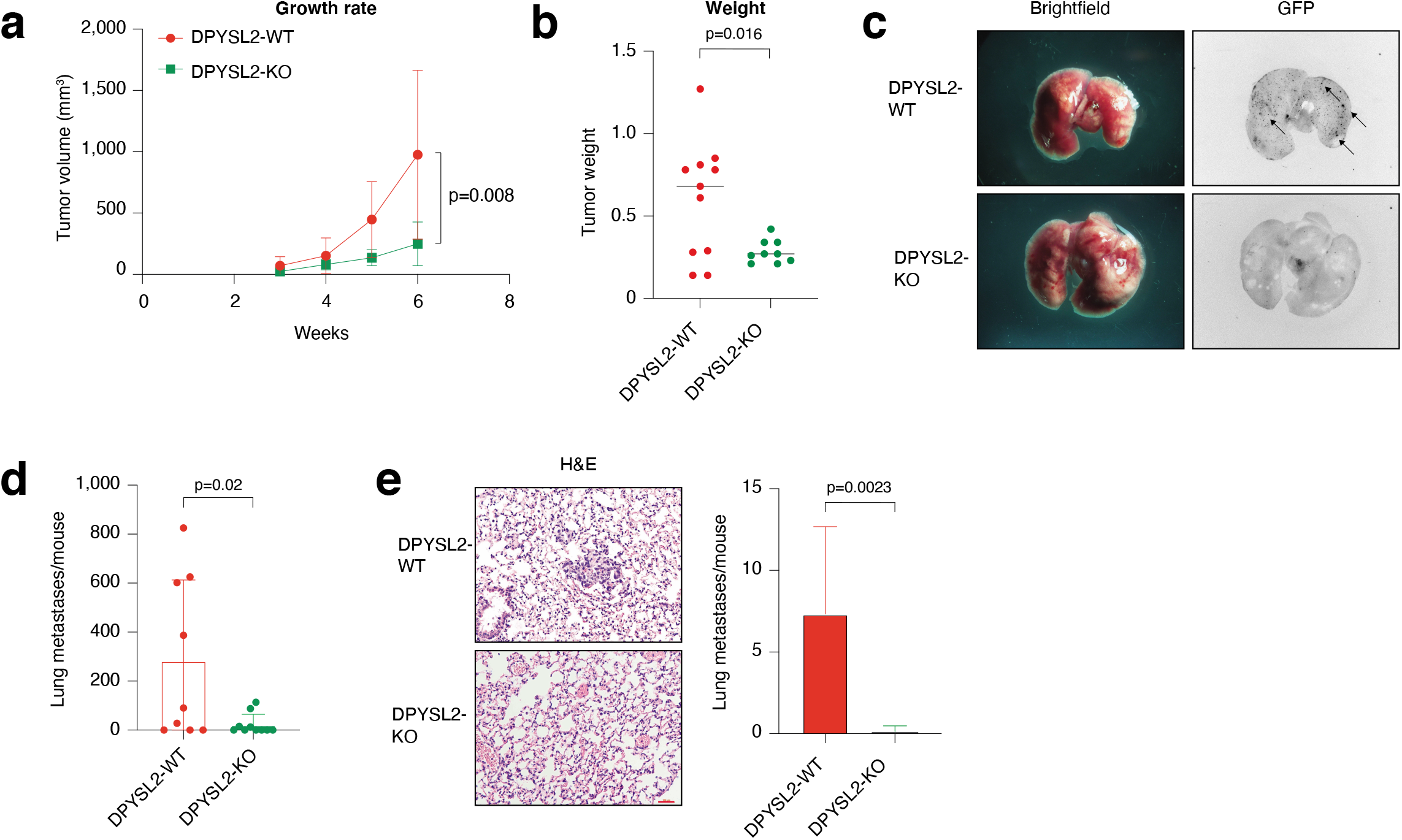
DPYSL2 loss affects tumor formation and metastasis in mice. (**a**) DPYSL2 expression affects tumor formation and growth rate in mice. MDA-MB-231 cells (WT or DPYSL2-KO) were injected into the mammary fat pad of female NOD-SGID mice. Over the indicated time course, both groups’ tumor volumes were measured and presented as a graph. Each value represents the mean ± SD. The p-value was determined by Student’s t-test, n = 11. (**b**) The tumors from (a) were weighed and presented as a graph. The p-value was determined by Student’s t-test, n = 11. (**c**) DPYSL2 loss reduces the number of lung metastases. Representative lungs obtained from WT-and DPYSL2-KO-injected mice. Left: brightfield images of the lung. Right: fuorescence images of GFP-labeled colonies. Arrows indicate the detected GFP-expressing metastases. (**d**) Quantification of the GFP-labeled lungs as described in (c). The p-value was determined by Student’s t-test. (**E**) DPYSL2 KO inhibits lung metastasis. Representative images (left) and quantification (right) of lung tissues stained with hematoxylin & eosin (H&E) from the same lungs in (c). The p-value was determined by Student’s t-test.

### JAK1 interacts with the C terminal domain of DPYSL2

DPYSL2 is a cytoskeleton-interacting protein that plays a vital role in axon guidance (46). This adaptor protein is regulated at its C-terminal domain (81 amino acids) (47) (Figure 4A) through posttranslational modifications (48). We performed an immunoprecipitation assay followed by LC-MS to identify DPYSL2-interacting partners in MDA-MB-231 cells, and then systematically categorized them using Metascape (Table S2). We found that the generated categories were associated with known DPYSL2 activities, such as folding of actin, axon guidance, and semaphorin interaction proteins (Figure S4A), which validates the capability of our assay. Interestingly, among the list of DPYSL2-interacting proteins, we identified JAK1, a crucial signaling molecule that mediates cancer cell aggressiveness (49). We found JAK1-DPYSL2 binding to be specific (DPYSL2-FLAG) since JAK1 did not interact with the control protein RAP2A (RAP2A-FLAG). In addition, this interaction is C-terminal dependent as JAK1 only binds the full-length protein and not the truncated form (DPYSL2-ΔCter-FLAG) (Figure 4B). This interaction was also observed in Hs-578-T cells, another basal B breast cancer cell line. To support these findings, we analyzed JAK1 expression in breast cancer patients and demonstrated a significant correlation with DPYSL2 (Figure 4C) and EMT markers (Figure S4B). In parallel, DPYSL2 levels significantly correlated with expression of cytokines and their receptors (Figure S4C), which are established targets of JAK1 (50). Together, these data show that DPYSL2 interaction is not exclusive to cytoskeletal proteins as it binds components of intracellular signaling pathways.

**Figure 4:**
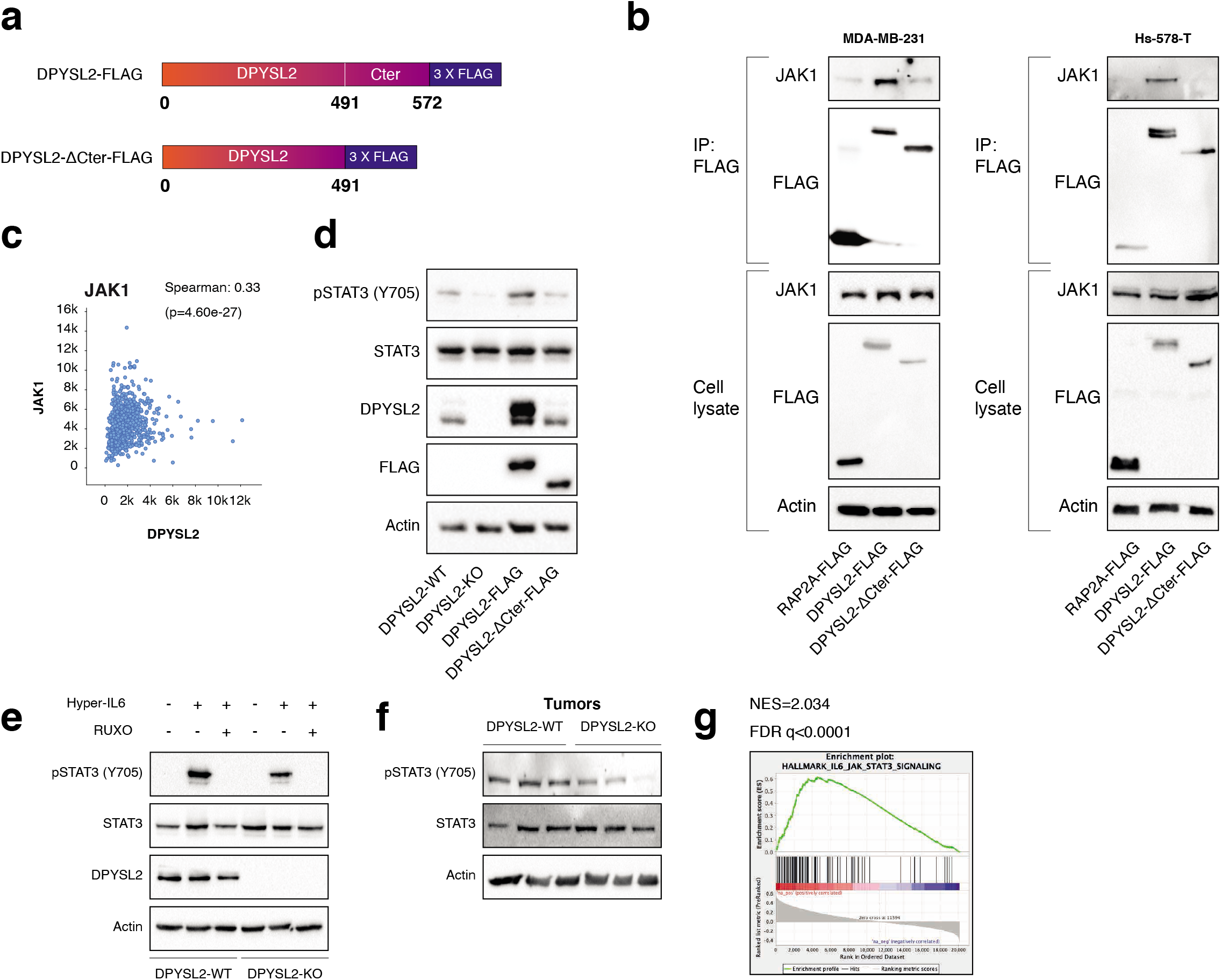
DPYSL2 interacts with JAK1 and regulates STAT3 signaling. (**a**) A schematic diagram of the amino acid alignment of full-length DPYSL2 and C terminal isoform. (**b**) JAK1 interacts with the regulatory C-terminal domain of DPYSL2. MDA-MB-231 cells expressing RAP2A-FLAG, DPYSL2-FLAG, and DPYSL2-Cter-FLAG were lysed and subjected to immunoprecipitation using anti-FLAG antibodies. Cell lysates and immunoprecipitates were analyzed by immunoblotting for the levels of indicated proteins. (**c**) DPYSL2 expression correlates with JAK1 in breast cancer patients. Patients’ gene expression data were generated by the TGGA project and analyzed using the cBioportal webtool (https://www.cbioportal.org). DPYSL2 expression positively and significantly correlated with JAK1. The Spearman correlation coefficient and the p-value were calculated by the analysis tool. (**d**) STAT3 phosphorylation correlates with DPYSL2 expression. Immunoblot representing DPYSL2 knockout (DPYSL2-KO) in MDA-MB-231, overexpression of the full-length variant (DPYSL2-FLAG) or the C-terminal truncated (DPYSL2-Cter-FLAG). Cells were lysed and subjected to immunoblotting using the indicated antibodies. The DPYSL2 antibody targets the C-terminal region, thus it does not detect DPYSL2-Cter-FLAG. (**e**) Loss of DPYSL2 expression results in STAT3 signaling inhibition. DPYSL2-WT and DPYSL2-KO cells were starved with 0% FBS medium for 16 h and treated with 0 and 15 µl of media from HEK-293 cells generating Hyper IL-6 for one hour. For the inhibitor samples, 30 mins before Hyper IL-6 treatment, cells were incubated with 30µM ruxolitinib (JAK1/2 inhibitor). Cells were subjected to immunoblot using the indicated antibodies. (**f**) Tumors developed from DPYSL2-KO cells showed a reduction in STAT3 phosphorylation. Three representative tumors originated from WT or DPYSL2-KO cells were lysed and subject to immunoblotting using the indicated antibodies. (**g**) DPYSL2 expression in breast cancer patients correlates with the hallmark of IL6/JAK/ STAT3 signaling. Breast cancer patients’ gene expression data were generated by the TGGA (PanGancer Atlas project) and analyzed using the cBioportal webtool (https://www.cbioportal.org). In these samples, the expression of DPYSL2 was compared to the whole transcriptome (∼20,000 genes). The genes were ranked based on the obtained Spearman rank-order correlation coefficient following by GSEA analysis. GSEA computed the NES and FDR values.

### DPYSL2 as a regulator of STAT3 signaling

Upon its activation, JAK1 phosphorylates the transcription factor STAT3 on tyrosine 705 (51), which in turn translocates into the nucleus and induces the expression of selected metastatic and migratory factors, such as vimentin and MMP9 (19). After identifying DPYSL2-JAK1 interaction, we determined whether this adaptor protein plays a role in STAT3 activation. Indeed, compared to WT cells, the basal level of STAT3 phosphorylation was decreased in knockout cells (DPYSL2-KO) and increased in cells ectopically expressing the full-length DPYSL2 (DPYSL2-FLAG) (Figure 4D). In contrast, overexpression of the truncated form (DPYSL2-ΔCter-FLAG) did not affect STAT3 phosphorylation, thus indicating the importance of the C-terminal domain in regulating this signaling cascade.

To further establish DPYSL2’s role in STAT3 signaling, we treated both WT and DPYSL2-KO cells with Hyper-IL6 (52), a fusion protein of IL-6 and the soluble IL6-receptor (sIL6R), which is a potent inducer of STAT3 phosphorylation (53). We found that Hyper-IL6 substantially induced STAT3 phosphorylation in DPYSL2-WT cells, while this induction was lower in knockout cells (Figure 4E and S4D). Moreover, we verified activation of the IL6/JAK/STAT3 axis by co-treating the cells with Hyper-IL6 and ruxolitinib, a selective inhibitor of JAK1 and JAK2 (54), which abolished the STAT3 phosphorylation. Similarly, xenografts generated from DPYSL2-WT cells exhibited higher STAT3 phosphorylation than those from DPYSL2-KO cells (Figure 4F). Finally, in breast cancer patients, DPYSL2 expression significantly correlated with IL-6/JAK/STAT3 signaling (Figure 4G). Together these findings indicate that DPYSL2 is a critical factor in the activation of the JAK1-STAT3 signaling cascade.

### DPYSL2 mediates STAT3 induction of vimentin expression

Our differential transcriptome analysis identified a group of 2,302 genes that are downregulated upon DPYSL2 loss (DPYSL2-DOWN). By analyzing the 48 STAT3 target genes’ distribution, we identified 16 that are significantly enriched in the DPYSL2-DOWN group (Figure S5A and S5B), including vimentin. Next, we assessed the vimentin protein level by immunoblot and immunofluorescence and found that DPYSL2-KO cells expressed less vimentin relative to WT (Figure 5A and S5C). Moreover, overexpression of the full-length DPYSL2 (DPYSL2-FLAG), but not the C-terminal truncated form (DPYSL2-ΔCter-FLAG), resulted in elevated vimentin expression level in knockout cells (Figure 5A). To further determine that DPYSL2 functions upstream of the STAT3-vimentin axis, we ectopically expressed the constitutively activated STAT3 (A662C, N664C, V667L, (CA-STAT3)) (55) in DPYSL2-KO cells. We found that in these cells, CA-STAT3 promoted vimentin expression (Figure 5B) and migration capability (Figure 5C and S5D). Furthermore, CA-STAT3 induced its known target IL-6 (51), which validated its activity (Figure S5E). Finally, we clarified DPYSL2’s role in cell migration, as its loss disrupted the actin-based structures known as filopodia at the edge of the migrating cells (Figure 5D). Together these results demonstrated that DPYSL2 governs cancer cell migration through its essential role in the JAK1/STAT3/vimentin axis.

**Figure 5:**
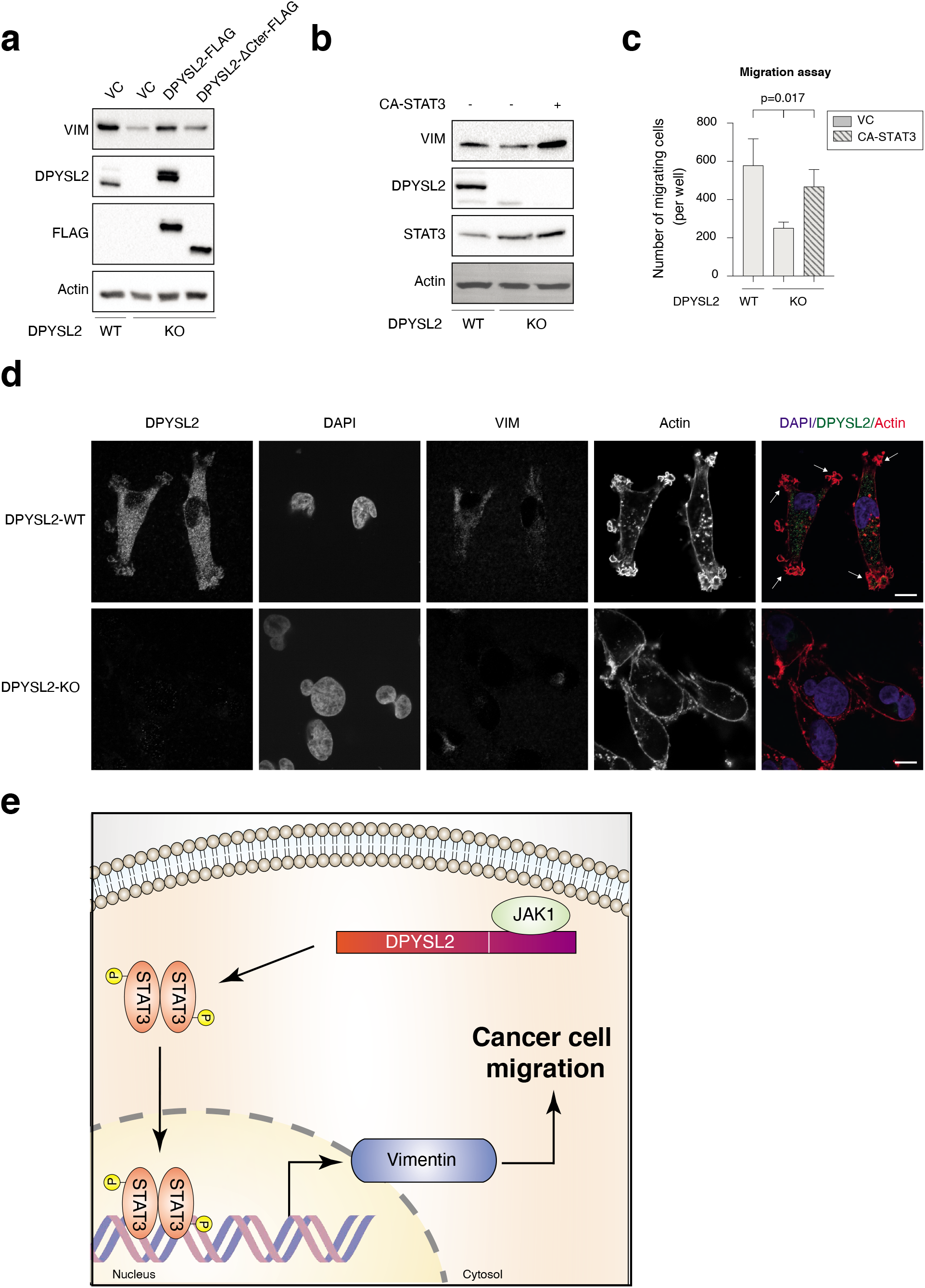
DPYSL2 regulates JAK/STAT3 signaling. (**a**) DPYSL2 expression correlates with vimentin expression. Immunoblot representing DPYSL2 knockout (DPYSL2-KO) in MDA-MB-231 cells, and overexpression of the full-length variant (DPYSL2-FLAG) or the C-terminal truncated (DPYSL2-Cter-FLAG) in the KO background. Cells were lysed and subjected to immunoblotting using the indicated antibodies. VC -vector control. VIM -vimentin (**b**) Constitutively activated STAT3 induces vimentin expression. Immunoblots representing DPYSL2 knockout (DPYSL2-KO) in MDA-MB-231 and overexpression of the constitutive activates STAT3 (CA-STAT3) in the KO background. Cells were lysed and subjected to immunoblotting using the indicated antibodies. Vim=vimentin. VIM -vimentin (**c**) Constitutively activated STAT3 (CA-STAT3) improves the migration of DPYSL2-KO cells. The same cells as in (b) were subjected to transwell assay. The data is reported as the number of migrated cells per 20,000 seeded cells; each value represents the mean ± SD for n=3. The p-value was determined by Student’s t-test. VG -vector control. (**d**) DPYSL2 loss affects the cellular distribution of actin flaments. WT and DPYSL2-KO cells were subjected to immunofuorescence imaging using the indicated antibodies. Phalloidin was used to probe actin. Arrows indicate flopodia. Bar=10 µm. VIM -vimentin (**e**) A scheme representing the role of DPYSL2 in JAK/STAT3/vimentin-mediated cell migration.

## Discussion

We established the axon guidance adaptor protein DPYSL2 as a key factor in breast cancer cell aggressiveness. We found a significant correlation between the expression of DPYSL2 and mesenchymal markers in breast cancer patients and in cell lines. Moreover, DPYSL2 loss had a substantial impact on cell motility, tumorigenesis, and metastasis. We demonstrated that DPYSL2 regulates cancer cell migration through its binding to JAK1. This interaction induces STAT3 signaling, which upregulates the expression of migratory factors such as vimentin (Figure 5E). These findings indicate a novel cellular regulatory mechanism, where adaptor proteins serve as a direct link between cytoskeletal molecules and oncogenic signaling kinases. In this study, we demonstrated that cancer cells utilize components from the neuronal axon guidance machinery, including DPYSL2, to migrate efficiently. However, despite the similarities between cancer cell migration and axon guidance, these cellular systems regulate DPYSL2 expression differently. For instance, during axon growth, DPYSL2 is expressed at a relatively constant rate, but undergoes a diverse set of posttranslational modifications, such as phosphorylation, glycosylation, and SUMOylation (7). Nevertheless, DPYSL2 demonstrates a differential gene expression pattern in various tumor types. In colorectal carcinoma and non-small-cell lung cancer (NSCLC), DPYSL2 expression is upregulated in tumors relative to surrounding normal tissues (56-58), whereas epithelial breast cancer samples exhibited the opposite expression pattern (59). These cancer types also differ in their DPYSL2 expression pattern throughout tumor progression. In colorectal carcinoma and NSCLC, DPYSL2 expression does not vary between low-and high-grade samples, while in lung cancer, it demonstrates a progression-dependent phosphorylation pattern (57). However, in breast cancer, DPYSL2 level is upregulated in highly aggressive subtypes including basal and mesenchymal-like (60). This reported pattern correlates with METABRIC and TCGA databases, as our analysis shows that high DPYSL2 is enriched in normal, basal, and claudin-low samples (Figure 1D). Hence, these expression variations among diverse tumors suggest that DPYSL2 serves a cancer type-dependent function that is yet to be resolved.

By applying the CRISPR-Cas9 knockout system, we found that DPYSL2 expression is essential for the breast cancer cells to maintain their high-grade characteristics. However, a recent study, in which DPYSL2 expression was manipulated by overexpression or short hairpin RNA-mediated knockdown, suggested that this gene suppresses breast cancer aggressiveness (60). Here, we suggest that a defined expression level of DPYSL2 is crucial for its efficiency, as any variation may interfere with its stoichiometric interaction with JAK1 and consequently affect cell migration. The different experimental tools used in the two studies resulted in different DPYSL2 expression levels, which could generate dissimilar outcomes.

In neuronal development, DPYSL2 guides axons through direct binding with numerous cytoskeletal proteins, including actin and tubulin (61). Here, we identified that in breast cancer, DPYSL2 interacts with signaling factors as an additional mechanism to modulate cell migration. Specifically, we found that this adaptor protein plays an essential role as a regulator of the JAK1/STAT3 signaling cascade and subsequently governs vimentin expression. Thus, we predict that future studies will establish a similar regulatory role of DPYSL2 in developing neurons’ signaling cascades.

In addition to their role in development, many axon guidance genes, including DPYSL2, function in the neuronal injury response machinery (62,63). However, the function of DPYSL2 in neuronal regeneration is still an enigma. Upon neuronal injury, a set of pro-regenerative factors are released, including the cytokine IL-6 (64), which activates the JAK/STAT3 signaling cascade (65). Since we determined that DPYSL2-JAK1 interaction is a regulator of STAT3 signaling in cancer cells, we suggest that the same mechanism applies to neuronal injury response. Thus, identifying factors that enhance DPYSL2-JAK1 interaction in cancer cells can be potentially used as drug targets to improve neuronal injury repair.

## Supporting information

CEL-Seq analysis of DPYSL2 -WT and KO cells

METASCAPE analysis of DPYSL2 interacting proteins

## Acknowledgments

We thank the members of the Y.D.S. laboratory. This work was supported by the Israel Science Foundation (Grant 1816/16), the Israel Cancer Research Fund RCDA fellowship, and the Hebrew University start-up funds. A.A.R. is supported by the Brodie fellowship for breast cancer research. B.S. is supported by the Lady Davis Fellowship for postdoctoral researchers at The Hebrew University of Jerusalem. Dr. William Breuer, Proteomics/Mass spectrometry Unit, Institute of Life Sciences, Hebrew University of Jerusalem, for his help in the LC-Ms experiments and analysis. Amina Jbara, Hebrew University, for assisting us with the animal experiments.

## Author contribution

A.A.R and Y.D.S designed the study. A.A.R performed most of the experiments. B.S. performed most of the *in vivo* studies. M.B.Y., together with A.A.R., performed many of the western blots and qPCR studies. A.K. generated the viruses. M.L. performed most of the cloning. M.T. performed many of the FACS studies. A.H., B.S. and A.A.R performed the *in vitro* functional assays. N.P. analyzed the IHC pictures. Y.D.S. and A.A.R. wrote the paper with input from all the authors.

