## Supplementary Data for "Dihydropyrimidinase-like 2 (DPYSL2) regulates breast cancer migration via a JAK/STAT3/vimentin axis"

***Supporting Figures***

**Figure S1**: DPYSL2 expression is elevated in mesenchymal-like cells. (**A**) A table representing the expression ratio of selected genes in epithelial and mesenchymal cell lines and its *p*-value. The *p*-value was determined by Student's *t*-test. (**B**) *DPYSL2* expression correlates with axon guidance genes. Patients' gene expression data were generated by the TCGA project and analyzed using the cBioportal webtool (https://www.cbioportal.org). DPYSL2 expression positively and significantly correlated with axon guidance genes (*ROBO1*, *SLIT2*, *FYN,* and *NRP1*). The Pearson's and Spearman's correlation coefficients, and the *p*-value, were calculated by the analysis tool. (**C**) DPYSL2 expression positively and significantly correlated with known mesenchymal markers (*VIM*, *SNAI2, ZEB1*, and *ZEB2*). (**D**) DPYSL2 expression negatively and significantly correlated with known epithelial markers (*KRT18* and KRT*19*).

**Figure S2:** DPYSL2 loss inhibits cell migration in breast cancer cell lines. **(A)** Representative pictures of the wound healing assay. Representative images of relative wound density at 0 and 24 h for the indicated WT, DPYSL2-KO, and DPYSL2-KO+ DPYSL2-FLAG cells. (**B**) DPYSL2 KO does not affect proliferation. The proliferation rate of MDA-MB-231 WT, DPYSL2-KO, and DPYSL2-KO+DPYSL2-FLAG cells were measured using CellTiter-Glo. N.S.= no statistically significant differences between the samples. (**C**) Loss of DPYSL2 results in CD44 cell surface expression reduction. The different indicated samples were subjected to FACS analysis of the cell-surface marker CD44. The histogram represents CD44 fluorescence intensity values. n=3.

**Figure S3:** DPYSL2 loss affects tumor formation and metastasis in mice. (**A**) Tumor formation in mice. Representative pictures of tumors taken out from mice injected with DPYSL2-WT cells (top) and DPYSL2-KO cells (bottom). (**B**) DPYSL2 loss reduces the number of lung metastases. Representative lungs obtained from WT- and DPYSL2-KO-injected mice. Left:

brightfield images of the lung. Right: fluorescence images of GFP-labeled colonies. Arrows indicate the detected GFP-expressing metastases. (**C**) DPYSL2 KO inhibits lung metastasis. Representative images of lung tissues stained with hematoxylin & eosin (H&E) from the same lungs in (B).

**Figure S4:** DPYSL2 interacts with JAK1 and regulates STAT3 signaling. (**A**) DPYSL2-FLAG interacts with axon guidance proteins. The functional categories of proteins interacting with DPYSL2 as identified by the Metascape analysis (1). (**B**) JAK1 expression in breast cancer patients correlated with the hallmark of EMT. Breast cancer patients' gene expression data were generated by TCGA (PanCancer Atlas project) and analyzed using the cBioportal webtool (2). In these samples, the expression of JAK1 was compared to the whole transcriptome (~20,000 genes). The genes were ranked based on the obtained Spearman's correlation coefficient followed by Gene set enrichment analysis (GSEA) analysis. GSEA computed the normalized enrichment score (NES) and FDR values. (**C**) DPYSL2 expression in breast cancer patients correlated with the KEGG cytokine and cytokine receptor interactions. Breast cancer patients' gene expression data was generated and analyzed as (B). (**D**) Loss of DPYSL2 expression results in STAT3 signaling inhibition. DPYSL2-WT and DPYSL2-KO cells were starved with 0% FBS medium for 16 h and treated for one hour with 0, 15, 25, 50, and 75 µl of media generated from HEK-293 cells generating Hyper IL-6. Cells were subjected to immunoblot using the indicated antibodies.

**Figure S5:** DPYSL2 regulates JAK/STAT3 signaling. (**A**) The expression of STAT3 target genes is enriched in the downregulated DPYSL2-KO gene set. Pie charts representing the proportion of STAT3-target genes in the significantly downregulated genes vs. unchanged ("STAT3 target genes") and all other genes ("all genes"). The enrichment of STAT3 target genes was quantified using a Fisher's exact test. (**B**) DPYSL2 loss induced a global reduction in the expression of STAT3 target genes. The expression levels in WT and DPYSL2 KO cells is presented in a volcano plot. STAT3 target genes are presented as red triangles. (**C**) DPYSL2 loss affected vimentin expression. WT and DPYSL2 cells were subjected to immunofluorescence imaging using the indicated antibodies. Bar=10µm. (**D**) CA-STAT3 rescued cell migration in the breast cancer cell line. Images representing the migratory capability of the different samples as determined by the transwell migration assay. The data is reported as the number of migrated cells per 20,000 seeded cells. (**E**). CA-STAT3 induced IL-6 expression in MDA-MB-231 cells. The RNA was isolated from WT, DPYSL2-KO, DPYSL2-KO+DPYSL2-FLAG, and DPYSL2-KO+CA-STAT3 cells, and the expression of the indicated genes was determined by qPCR. Each value represents the mean ± SD for n = 3. VC - vector control.

***Materials and methods***

**Cell Lines and Cell Culture**

The cell lines ZR-75-1, EVSA-t, MCF-7, MDA-MB-436, MDA-MB-231 and Hs-578-T were obtained from ATCC. ZR-75-1 cells were maintained in RPMI supplemented with 10% heat inactivated fetal bovine serum (FBS), while the remaining cell lines were maintained in DMEM (10% FBS). The immortalized human mammary epithelial cells expressing OHT-inducible Twist (HMLE-Twist-ER), and Naturally Arising MEsenchymal Cells (NAMEC) have been described (3,4) and (5), respectively. HMLE-Twist-ER and NAMEC cells were maintained in MEGM (Lonza) growth media. All cells were cultured at 37°C with 5% $CO_2$. For EMT induction, HMLE-Twist-ER cells were treated with 4-hydroxytamoxifen (OHT) (Sigma-Aldrich) at a final concentration of 10 nM for the indicated number of days.

**Cell Lysis and Immunoblotting**

Cells were rinsed twice with ice-cold PBS and lysed with RIPA lysis buffer (20 mM Tris pH 7.4, 137 mM NaCl, 10% glycerol, 1% Triton X-100, 0.5% deoxycholate, 0.1% SDS, 2.0 mM ethylenediaminetetraacetic acid (EDTA, pH 8.0), EDTA-free protease inhibitor (1 tablet per

50 ml RIPA) (Roche) and phosphatase inhibitor cocktail (100X) (Bimake). The lysates were cleared by centrifugation at 13,000 rpm at 4°C in a microcentrifuge for 10 min. The protein concentration was determined by Bradford (BioRad). Proteins were denatured by the addition of SDS sample buffer (5X) and boiling for 5 min, then resolved by 10% SDS-PAGE, transferred onto a 0.45 µm PVDF membrane (Merck), and probed with the appropriate antibodies.

**Antibodies**

Antibodies were obtained from the following sources: JAK1 (133666) and vimentin (ab8069) from Abcam; DPYSL2 (35672), CDH1 (3195), β-actin (4970)/(3700), vimentin (5741), FLAG (8146), p-STAT3-Tyr705 (9145) and STAT3 (9139) from Cell Signaling Technology; APC-labeled anti-CD44 (559942) from BD Bioscience; HRP-labeled anti-mouse (Jackson ImmunoResearch Laboratories, 115-035-003) and anti-rabbit (111-035-144) secondary antibodies from Jackson ImmunoResearch. For immunofluorescence assays: Goat-anti mouse-Alexa fluor 647 (Abcam, ab150119), Donkey-anti-rabbit-Rhodamine red X (Jackson ImmunoResearch Laboratories, 711-295-152) and donkey anti rabbit-alexa fluor 488 (Jackson ImmunoResearch Laboratories, 711-545-152)

**Cancer Sample Analysis**

KM analysis of the data from breast cancer samples were analyzed and generated by the KM Plotter website (6),(7). Search entries: DPYSL2 as the gene symbol, (Affymetrix ID: 200762_at), auto-select best cutoff for "split patients by." The obtained KM plots and the statistics were generated by the website.

**Plasmids**

DPYSL2 was received from the CCSB human orfeome collection hORFeome V7.1 ID12017 http://horfdb.dfci.harvard.edu/hv7/index.php?page=getresults&by=detail&qury=12017 and cloned into pLJC2 (8) kindly provided by Dr. Susan Lindquist . DPYSL2-ΔCter-FLAG was cloned by us. STAT3 ((A662C, N664C, V667L)-pcw107-V5 was a gift from David Sabatini & Kris Wood (Addgene plasmid # 64611; http://n2t.net/addgene:64611; RRID: Addgene_64611). pLJC2-Rap2A-3xFLAG was a gift from David Sabatini (Addgene plasmid # 87974 ; http://n2t.net/addgene:87974 ; RRID:Addgene_87974).

**Microscopy**

The MDA-MB- 231 ($6 \times 10^4$) cells were seeded on polylysine-coated glass coverslips in 12-well tissue culture plates. After 24 h, the slides were rinsed twice with PBS and fixed with 4% paraformaldehyde in PBS for 15 min at room temperature, followed by quenching with ammonium chloride (1% in PBS). The slides were then rinsed three times with PBS, and cells were permeabilized with 0.1% Triton X-100 in PBS for 10 min. The cells were gently rinsed three times with PBS and subsequently incubated for 30 mins in the blocking buffer (1% BSA in PBST), followed by primary antibodies for 1 h (DPYSL2 and vimentin 1:200 each in 1%BSA in TBST). The cells were rinsed three times with PBS, incubated with secondary antibodies (diluted 1:200 in 1% BSA in TBST) and Phalloidin-iFluor 555 Reagent (Abcam 1:5000) for 1 h at room temperature in the dark, and washed three times PBS. Slides were mounted on glass coverslips using Vectashield (Vector Laboratories) and cells imaged on a spinning disk confocal system (Nikon).

**Virus Production**

HEK-293T cells were co-transfected with the pLentiCRISPR sgRNA, VSV-G envelope plasmid, and Δvpr lentiviral plasmid using X-TremeGene 9 Transfection Reagent. The supernatant containing the virus was collected 48 h after transfection and spun for 5 min at 400 g to eliminate cells.

**Cell Proliferation Assay**

Cells were seeded in white 96-well plates (Greiner) at a density of $0.5 \times 10^3$ cells/well (MDA-MB 231 WT, DPYSL2 KO, DPYL2-KO+DPYSL2-FLAG). Cell viability was assessed with

Cell Titer-Glo (Promega) after 1-, 3-, and 5-days following seeding, and luminescence was measured with Cytation 3 (Biotek).

**FACS Analysis**

MDA-MB-231 ($2 \times 10^5$) cells were seeded in a 6-well tissue culture plate. After 16 h, cells were washed 3 times with PBS and detached for further processing using 200 µl of 0.05% EDTA solution. Further, cells were quenched using 10 % FBS DMEM (Biological industries), collected in Eppendorf tubes, centrifuged (2,500 rpm for 5 mins) at 4 °C, then washed with PBS twice. Next, cells were incubated with CD44 APC conjugated antibody for 30 mins. The cells were then washed with PBS twice and were passed through filter mesh and collected in FACS tubes. Samples were sorted by the flow cytometer BD Accuri C6 and analyzed by FlowJo software (Tree Star).

**Mammosphere Formation Assay**

Eight hundred cells/well were seeded in 96-well plates after coating with growth-factor-reduced Matrigel (Corning). Plates were incubated for 5-8 days under observation, and every two days, 2% of new growth-factor-reduced Matrigel was added. Mammospheres were counted manually under the microscope and plotted as a graph.

**Wound Healing Assay**

MDA-MB-231 cells ($4 \times 10^4$) were plated on to IncuCyte ImageLock 96-well cell culture microplates. After 18 h, the cell monolayer was scraped using a wound-maker mechanical device (Essen BioScience), washed with PBS, and examined under an inverted microscope. The wound area was monitored using the IncuCyte live-cell imaging system. Wound healing assay results were compiled from eight wells with one scratch in each well. At the 24 h time point, closure of the control scratch was observed.

**Transwell Migration and Invasion Assays**

Migration assays were performed using the Costar Transwell Invasion chamber (USA). Transwell inserts were hydrated with serum-free DMEM for 30 mins, and 2 x $10^4$ of MDA-MB-231 cells (500 µl) were added to the upper chamber. The invasion assay was performed using Millicell cell culture inserts (Millipore). The inserts were coated with Extracellular Matrix (ECM, 1ug/ml) with serum-free DMEM for 1 h, and 2 × $10^4$ of MDA-MB-231 cells (500 µl) were added to the upper chamber. For both assays, DMEM (700 µL) with 10% FBS (as chemoattractant) was added to the lower wells of the 24-well plate. The medium was discarded after 24 h. Non-migratory cells were removed with cotton-tipped swabs, and the lower surface of the insert was stained with 0.5% crystal fast violet. The cells were counted and captured under a Nikon Eclipse 80i microscope at 10× magnification.

**RNA Preparation, RT-PCR and qPCR Analysis**

| Name | Forward | Reverse |
|---|---|---|
| IL-6 | ACTCACCTCTTCAGAACGAATTG | CCATCTTTGGAAGGTTCAGGTTG |
| GAPDH | AGCCACATCGCTCAGACAC | GCCCAATACGACCAAATCC |
| DPYSL2 | GATCCCCGGAGGAATTGACG | GGCTCAGGAACAACGTGGTC |

Total RNA was isolated from cells using the NucleoSpin® RNA Kit (MACHEREY-NAGEL, Germany), and reverse-transcription was performed using qScript cDNA Synthesis Kit (Quantbio, USA). The resulting cDNA was diluted in DNase-free water (1:10) before quantification by real-time quantitative PCR. The mRNA transcription levels were measured using 2x qPCRBIO SyGreen. Blue Mix Hi-ROX (PCR Biosystems) and StepOnePlus (Applied Biosystems). All data were expressed as the ratio between the expression level of the target gene mRNA and GAPDH. The list of primers used for the RNA-Seq analysis were obtained from Integrated DNA Technology and are listed below:

**Hematoxylin and Eosin Staining**

Lung tissue sections were stained with H&E using Sakura Tissue-Tek Prisma (Department of Pathology, Hadassah Hebrew University Medical Center). Lung metastasis incidence was analyzed by a pathologist (NP). Quantification of lung metastatic load was performed by analyzing the number and volume of metastatic lesions per section.

***Supplementary Tables***

**Table S1:** CEL-Seq analysis of DPYSL2 -WT and KO cells.

**Table S2:** METASCAPE analysis of DPYSL2 interacting proteins.

***Supplementary References***

a

| Gene name | Ratio | p value |
|---|---|---|
| ROBO1 | 1.4828 | 9.5e-12 |
| SLIT2 | 1.4531 | 1.56e-10 |
| NRP1 | 1.4217 | 5.19e-19 |
| DPYSL2 | 1.418 | 1.58e-18 |
| CFL2 | 1.4 | 1.02e-19 |
| FYN | 1.3421 | 4.73e-18 |
| CXCL12 | 1.3007 | 2.59e-08 |
| EPHA5 | 1.1122 | 2.077e-07 |
| GNA12 | 1.0881 | 9.439e-28 |
| SAMA5A | 1.0084 | 7.638e-09 |
| EPHA1 | -1.0984 | 1.246e-47 |
| PAK6 | -1.153 | 1.062e-47 |
| RHOD | -1.2004 | 3.51e-42 |
| ABLIM1 | -1.4718 | 3.186e-43 |
| EFNA1 | -1.9129 | 1.56e-22 |

b

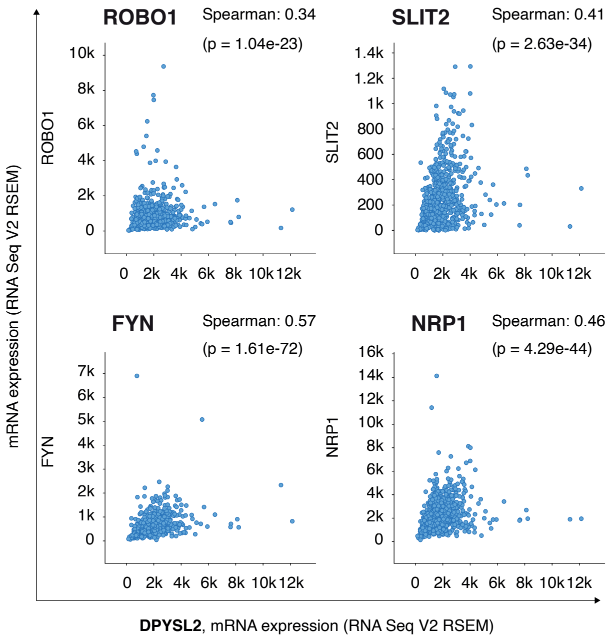


c

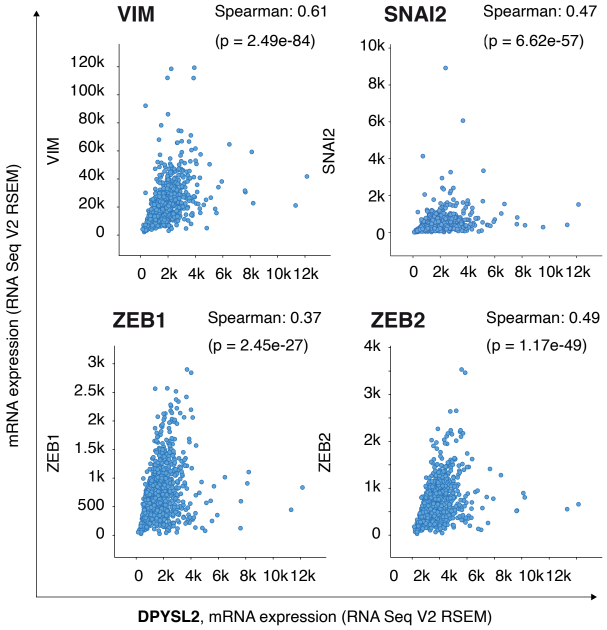


d

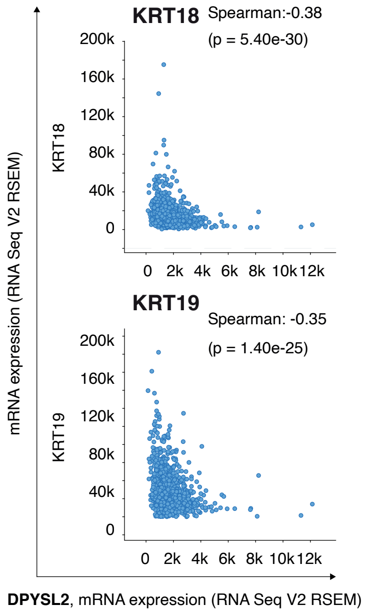


**Figure S1: DPYSL2 expression is elevated in mesenchymal-like cells.** (**a**) A table representing the expression ratio of selected genes in epithelial and mesenchymal cell lines and its p-value. The p-value was determined by Student's t-test. (**b**) DPYSL2 expression correlates with axon guidance genes. Patients' gene expression data were generated by the TCGA project and analyzed using the cBioportal webtool (https://www.cbioportal.org). DPYSL2 expression positively and significantly correlated with axon guidance genes (ROBO1, SLIT2, FYN, and NRP1). The Pearson's and Spearman's correlation coefficients, and the p-value, were calculated by the analysis tool. (**c**) DPYSL2 expression positively and significantly correlated with known mesenchymal markers (VIM, SNAI2, ZEB1, and ZEB2). (**d**) DPYSL2 expression negatively and significantly correlated with known epithelial markers (KRT18 and KRT19).

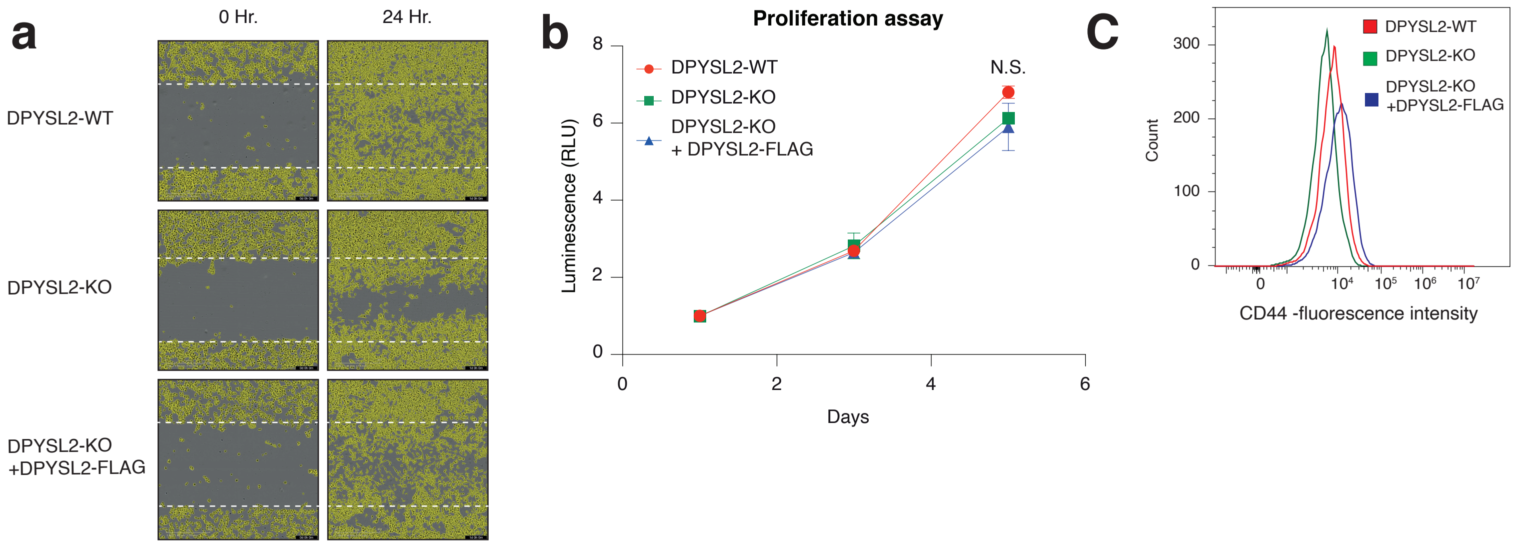


**Figure S2: DPYSL2 loss inhibits cell migration in breast cancer cell lines.** (**a**) Representative pictures of the wound healing assay. Representative images of relative wound density at 0 and 24 h for the indicated WT, DPYSL2-KO, and DPYSL2-KO+ DPYSL2-FLAG cells. (**b**) DPYSL2 KO does not affect proliferation. The proliferation rate of MDA-MB-231 WT, DPYSL2-KO, and DPYSL2-KO+DPYSL2-FLAG cells were measured using CellTiter-Glo. N.S.= no statistically significant differences between the samples. (**c**) Loss of DPYSL2 results in CD44 cell surface expression reduction. The different indicated samples were subjected to FACS analysis of the cell-surface marker CD44. The histogram represents CD44 fluorescence intensity values. n=3.

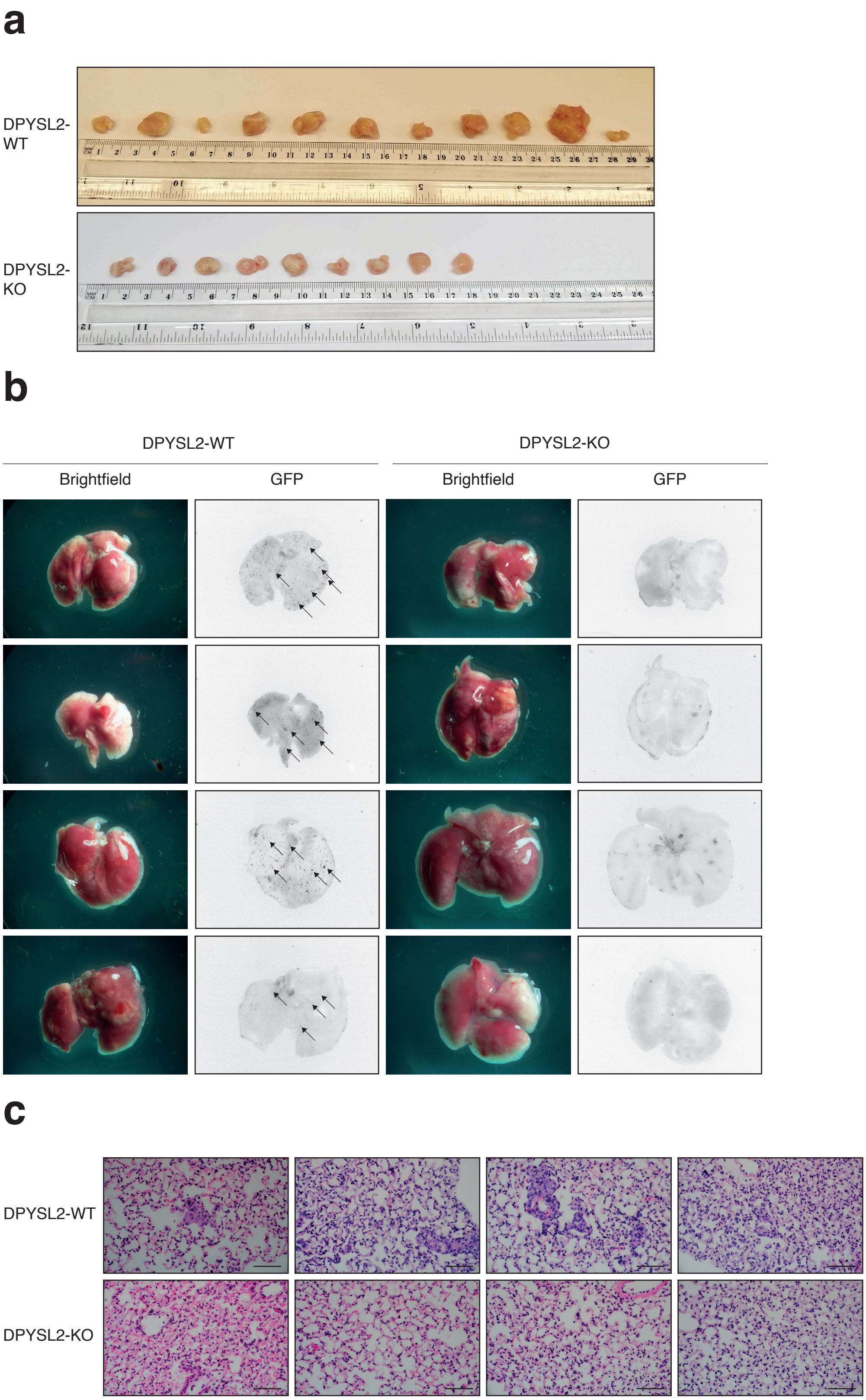


**Figure S3: DPYSL2 loss affects tumor formation and metastasis in mice.** (**a**) Tumor formation in mice. Representative pictures of tumors taken out from mice injected with DPYSL2-WT cells (top) and DPYSL2-KO cells (bottom). (**b**) DPYSL2 loss reduces the number of lung metastases. Representative lungs obtained from WT- and DPYSL2-KO-injected mice. Left: brightfield images of the lung. Right: fluorescence images of GFP-labeled colonies. Arrows indicate the detected GFP-expressing metastases. (**c**) DPYSL2 KO inhibits lung metastasis. Representative images of lung tissues stained with hematoxylin & eosin (H&E) from the same lungs in (b).

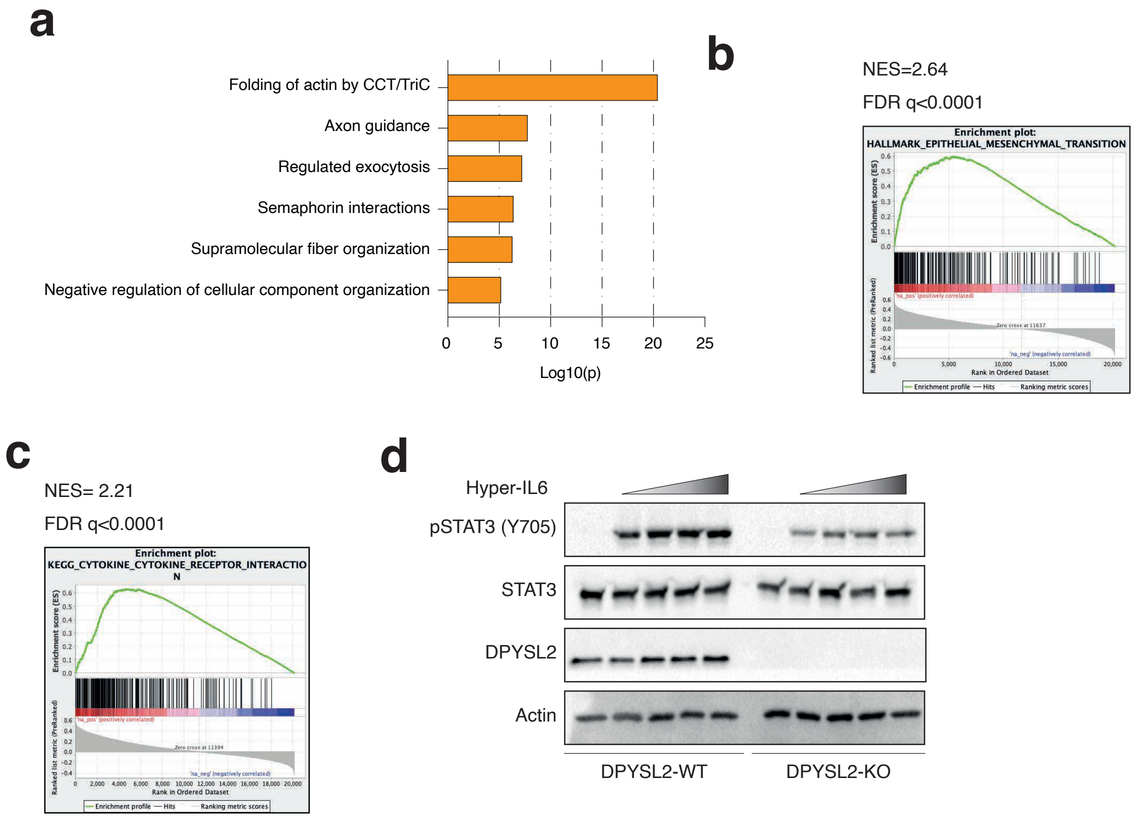


**Figure S4: DPYSL2 interacts with JAK1 and regulates STAT3 signaling.** (**a**) DPYSL2-FLAG interacts with axon guidance proteins. The functional categories of proteins interacting with DPYSL2 as identified by the Metascape analysis 1. (**b**) JAK1 expression in breast cancer patients correlated with the hallmark of EMT. Breast cancer patients' gene expression data were generated by TCGA (PanCancer Atlas project) and analyzed using the cBioportal webtool 2. In these samples, the expression of JAK1 was compared to the whole transcriptome (~20,000 genes). The genes were ranked based on the obtained Spearman's correlation coefficient followed by Gene set enrichment analysis (GSEA) analysis. GSEA computed the normalized enrichment score (NES) and FDR values. (**c**) DPYSL2 expression in breast cancer patients correlated with the KEGG cytokine and cytokine receptor interactions. Breast cancer patients' gene expression data was generated and analyzed as (B). (**d**) Loss of DPYSL2 expression results in STAT3 signaling inhibition. DPYSL2-WT and DPYSL2-KO cells were starved with 0% FBS medium for 16 h and treated for one hour with 0, 15, 25, 50, and 75 $\mu$l of media generated from HEK-293 cells generating Hyper IL-6. Cells were subjected to immunoblot using the indicated antibodies.

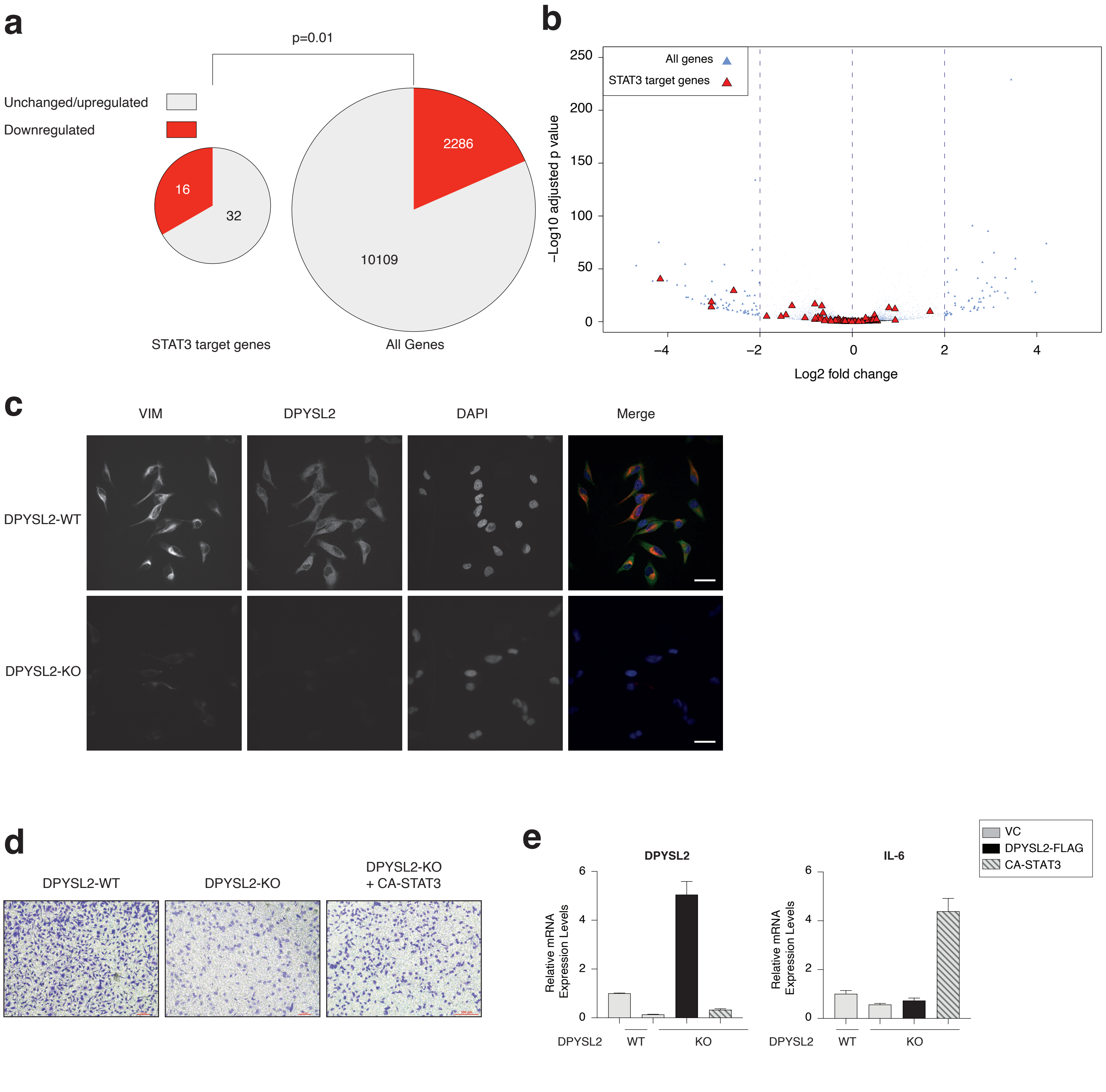


**Figure S5: DPYSL2 regulates JAK/STAT3 signaling.** (**a**) The expression of STAT3 target genes is enriched in the downregulated DPYSL2-KO gene set. Pie charts representing the proportion of STAT3-target genes in the significantly downregulated genes vs. unchanged ("STAT3 target genes") and all other genes ("all genes"). The enrichment of STAT3 target genes was quantified using a Fisher's exact test. (**b**) DPYSL2 loss induced a global reduction in the expression of STAT3 target genes. The expression levels in WT and DPYSL2 KO cells is presented in a volcano plot. STAT3 target genes are presented as red triangles. (**c**) DPYSL2 loss affected vimentin expression. WT and DPYSL2 cells were subjected to immunofluorescence imaging using the indicated antibodies. Bar=10$\mu$m. (**d**) CA-STAT3 rescued cell migration in the breast cancer cell line. Images representing the migratory capability of the different samples as determined by the transwell migration assay. The data is reported as the number of migrated cells per 20,000 seeded cells. (**e**). CA-STAT3 induced IL-6 expression in MDA-MB-231 cells. The RNA was isolated from WT, DPYSL2-KO, DPYSL2-KO+DPYSL2-FLAG, and DPYSL2-KO+CA-STAT3 cells, and the expression of the indicated genes was determined by qPCR. Each value represents the mean ± SD for n = 3. VC - vector control.
